# LongStitch: High-quality genome assembly correction and scaffolding using long reads

**DOI:** 10.1101/2021.06.17.448848

**Authors:** Lauren Coombe, Janet X Li, Theodora Lo, Johnathan Wong, Vladimir Nikolic, René L Warren, Inanc Birol

## Abstract

**Background:** Generating high-quality *de novo* genome assemblies is foundational to the genomics study of model and non-model organisms. In recent years, long-read sequencing has greatly benefited genome assembly and scaffolding, a process by which assembled sequences are ordered and oriented through the use of long-range information. Long reads are better able to span repetitive genomic regions compared to short reads, and thus have tremendous utility for resolving problematic regions and helping generate more complete draft assemblies. Here, we present LongStitch, a scalable pipeline that corrects and scaffolds draft genome assemblies exclusively using long reads.

**Results:** LongStitch incorporates multiple tools developed by our group and runs in up to three stages, which includes initial assembly correction (Tigmint-long), followed by two incremental scaffolding stages (ntLink and ARKS-long). Tigmint-long and ARKS-long are misassembly correction and scaffolding utilities, respectively, previously developed for linked reads, that we adapted for long reads. Here, we describe the LongStitch pipeline and introduce our new long-read scaffolder, ntLink, which utilizes lightweight minimizer mappings to join contigs. LongStitch was tested on short and long-read assemblies of three different human individuals using corresponding nanopore long-read data, and improves the contiguity of each assembly from 2.0-fold up to 304.6-fold (as measured by NGA50 length). Furthermore, LongStitch generates more contiguous and correct assemblies compared to state-of-the-art long-read scaffolder LRScaf in most tests, and consistently runs in under five hours using less than 23GB of RAM.

**Conclusions:** Due to its effectiveness and efficiency in improving draft assemblies using long reads, we expect LongStitch to benefit a wide variety of *de novo* genome assembly projects. The LongStitch pipeline is freely available at https://github.com/bcgsc/longstitch.

## Background

With the growing availability and accessibility of many different DNA sequencing technologies, generating high-quality *de novo* genome assemblies remains a crucial step in gaining important biological insights from the raw sequencing data. Constructing these *de novo* assemblies can enable a multitude of research aims, such as cancer genomics studies, analysis of non-model organisms, and population studies, to name a few. However, the complex and repetitive nature of genomes has long been a challenge in routinely achieving chromosome-scale genome assemblies[1].

To address these challenges, numerous developments in sequencing technologies have emerged. Many of these platforms provide long-range information to help resolve the problematic repeats, including linked reads[2, 3], optical maps[4], Hi-C data[5] and long reads[6]. These sequencing advances in turn inspire the development of new bioinformatics tools tailored to the specific characteristics of each data type, particularly in the genome assembly domain[1].

Long-read sequencing from Oxford Nanopore Technologies Ltd. (ONT, Oxford UK) and Pacific Biosciences of California, Inc. (PacBio) can generate reads in the kilobases up to the megabase range, a stark contrast to short-read sequencing, which generally produces 150-300bp reads. The long reads thus provide a rich resource of long-range genomic information, a feature that has proven extremely useful for genome assembly work [7]. While the error rates of long reads remain higher than typical short-read technologies such as those generated on the Illumina sequencing instrument, the read accuracy is improving with each new pore chemistry and advances in base-calling algorithms, and read accuracies now average between 87-98%[6, 8]. Furthermore, the throughput and cost of long-read sequencing is also improving, making it an increasingly competitive and accessible technology for many research groups and applications.

While recently developed long-read *de novo* genome assembly tools are generating assemblies that are highly contiguous, the sequences often harbour errors and underutilize long-range information. Therefore, these assemblies stand to gain from misassembly correction and further scaffolding, to maximize the rich genomic information provided by long reads and produce a more optimal solution [9, 10]. These assembly improvements can be vital to many downstream applications such as the analyses of regulatory elements, structural variations and gene clusters.

A number of genome scaffolders have been developed to contiguate draft genome assemblies using the information provided by long reads. These tools include LINKS[11], npScarf[12], OPERA-LG[13], SSPACE-LongRead[14], and, more recently, LRScaf[15]. Most of these tools utilize alignments of long reads to the draft assembly to infer joins between sequences, with the exception of LINKS, which uses a paired word of length *k* (*k*-mer) matching approach. LRScaf is the most recently developed long-read scaffolding tool, and generates a scaffold graph using long-read alignments from minimap2[16] or BLASR[17]. The graph is then manipulated in various ways, including edge filtering, transitive edge reduction and tip removal followed by a bi-directional traversal of the graph based on unique and divergent nodes to produce the final scaffolds.

In addition to the scaffolding stage being crucial for maximizing assembly contiguity, a preceding misassembly correction step is also important to achieve the highest quality final assembly, without which scaffolding-only utilities risk propagating structural errors. Effectively, when these errors are identified and rectified before scaffolding, the sequences then have the potential to be joined with the correct adjacent sequences. This step can therefore lead to both more correct and contiguous assemblies, as demonstrated by our linked-read assembly correction utility, Tigmint[18]. While there are multiple tools for correcting raw long reads directly[19], there are none that perform assembly correction using only long reads *de novo*.

Tools such as Pilon[20] and REAPR[21] perform misassembly correction using short reads, and others such as misFinder[22] additionally use a closely-related reference genome. While ReMILO[23] can use long reads in its assembly correction pipeline, it does not use this evidence exclusively, but in addition to short-read data and a reference genome.

A wide range of bioinformatics tools in various domains are based on the use of *k*-mers, but storing and operating on all *k*-mers in the input sequences can be very computationally expensive, especially for larger genomes. LINKS addresses this limitation by thinning the *k*-mer input set, an early form of sequence “minimizers”. The use of minimizer sketches has gained popularity in recent years, where only a particular subset of *k*-mers from the input sequences are considered, resulting in significant savings in runtime and memory usage[24]. Recently, we used this concept of minimizer sketches in our minimizer graph-based reference-guided scaffolder ntJoin[25].

Here, we present LongStitch, an efficient pipeline that corrects and scaffolds draft genome assemblies using long reads. LongStitch incorporates multiple tools developed by our group: Tigmint-long, ntLink, and optionally, ARKS-long. Tigmint-long and ARKS-long are misassembly correction and scaffolding utilities, respectively, previously developed for linked reads[18, 26, 27], and are now adapted to use long reads. Within LongStitch, we introduce our new long-read scaffolder, ntLink, which utilizes lightweight minimizer mappings to join contigs. We show that these tools used together in the LongStitch pipeline produce high-quality and contiguous assemblies that harbour fewer misassemblies compared to the current state-of-the-art.

## Implementation

### The LongStitch pipeline

LongStitch is implemented as a pipeline using a Makefile, and consists of assembly correction and scaffolding stages using long reads (Fig. 1a). The inputs to the pipeline are any draft sequence assembly and a set of matching long-read sequences. Genome assembly utilities in the LongStitch pipeline use the long-read data to improve upon the draft assembly to output a final, scaffolded genome assembly. The input assembly can be generated using any method and any data type, including the same long reads that are supplied to LongStitch. The first step of LongStitch is Tigmint-long, which identifies and breaks the input assembly at putative misassemblies. Then, our newly developed long-read scaffolder ntLink joins the assembly-corrected contigs together based on the long-range information. Optionally, an additional round of scaffolding can be performed using ARKS-long.

**Fig. 1.**
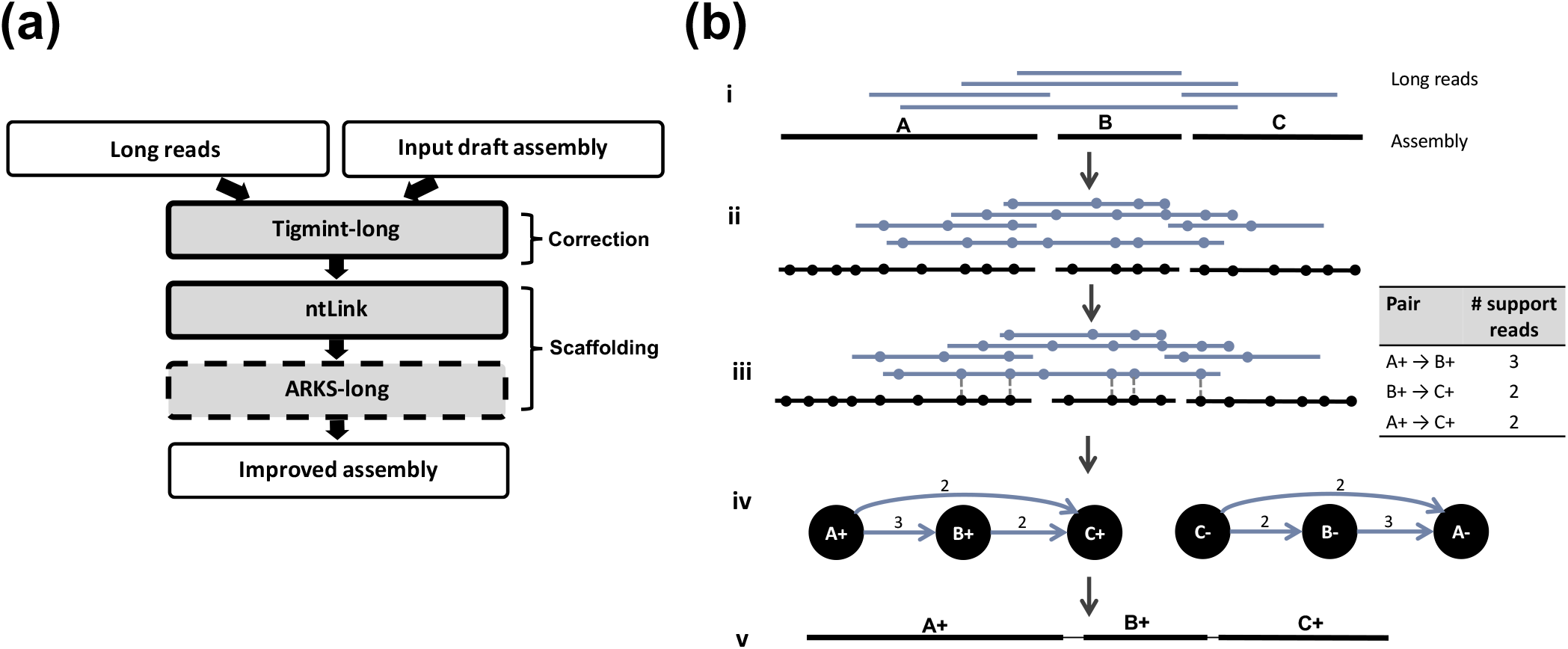
The LongStitch Pipeline. (a) Overview of the main steps of the LongStitch pipeline. The dashed border indicates the optional scaffolding step of ARKS-long. (b) Detailed schematic of the ntLink algorithm. (i) The input files to the ntLink long-read scaffolder are a draft assembly and long reads. (ii, iii) Minimizer sketches are computed for each of the input sequences (indicated by filled circles), then these minimizers are used to map the long reads to the draft assembly. Identical minimizers are vertically aligned, and also indicated by the vertical dotted line for one long-read. Each of the long-read mappings to the assembly can provide evidence about which contigs should be joined together, and in which orientation. (iv) After tallying all of the pairing information from the long reads, a directed scaffold graph is constructed, where the nodes are oriented contigs and the directed edges indicate long-read evidence between contigs. (v) Finally, the graph is traversed using abyss-scaffold to produce the final ordered and oriented scaffolds.

### Tigmint-long and ARKS-long

Tigmint[18] and ARKS[26, 27] are both previously published tools from our group. The tools were originally developed to correct and scaffold assemblies, respectively, both using linked reads. For LongStitch, we adapted these tools to use long reads as input by first generating pseudo-linked reads from the long reads. Briefly, tiled fragments are extracted from the long reads, and paired-end reads (each of length *fragment_size/2*, default *fragment_size*=500bp) are generated from the fragments. Each read pair extracted from the same long-read is assigned the same barcode, thus creating a reads file (in fastq or fasta) formatted like a linked reads file. These pseudo-linked reads are then input to the previously developed algorithms, which use the long-range information to perform correction and scaffolding.

In addition to generating the pseudo-linked reads, Tigmint and ARKS required other minor adjustments to work optimally with the long-read data. When using traditional linked reads, Tigmint uses bwa mem[28] to align the reads to the contigs, and then uses the alignments to infer molecule extents. The inferred molecule extents are examined to find and subsequently cut areas of the assembly that are not supported by these long molecules. In adapting Tigmint to work with long reads, we use minimap2[16] instead of bwa mem, which is better suited to long-read mappings. To reduce the number of parameters required for users to specify, we also added functionality to automatically calculate appropriate values for two parameters: *span* and *dist. Span* is the number of molecule extents spanning a region of the contig required for that region to be considered correct, and is set as: *span = 0*.*25 x long_read_coverage. Dist* is the maximum distance (in bp) allowed between read alignments of the same barcode for these reads to be merged to the same molecule extent, and is set as the median read length of the first one million long reads. The optimal values for both parameters are different for long reads compared to linked reads.

We also found that the optimal parameter settings for running ARKS-long varied from the recommended settings when using traditional linked reads. Particularly, using a smaller *k*-value (*k*=20), and a low Jaccard index threshold (*j*=0.05) was important to produce optimal scaffolding.

### ntLink

While the Tigmint-long and ARKS-long components of the pipeline are adaptations of previously published algorithms [18, 26, 27], within LongStitch we also introduce a new scaffolding tool, ntLink, which performs efficient long-read scaffolding using minimizer mappings (Fig. 1b).

First, minimizers are generated from both the input draft assembly and long reads as described in Roberts et *al*. [24], using a sliding window (*w*), and *k*-mer size (*k*) (Fig. 1b-i). Briefly, starting at the beginning of each sequence, the canonical hash values of *w* adjacent *k*-mers are generated using ntHash[29], and the smallest value is chosen as the minimizer for that window. For each minimizer, we also keep track of the corresponding sequence ID, the position (in bp) where the minimizer was found, and the strand of the canonical *k-*mer (“+” if the canonical *k*-mer is from the forward strand, else “-”). When applied over all of the input data, this process generates an ordered minimizer sketch for all input sequences.

Then, these minimizer sketches are used to map the long reads to the input draft assembly (Fig. 1b-ii). Any minimizers that are not unique in the draft assembly are discarded to avoid ambiguous mappings due to repeats. For each minimizer in a long read’s ordered sketch, the minimizer sketch of the draft contigs is queried. For every minimizer match in the draft contigs sketch, the information about which contig the minimizer hits to, as well as the position and strand of that minimizer in the contig is readily available. By performing this matching for each minimizer in the ordered sketch for a long read, we convert the minimizer sketch to an ordered list of contig hits (Ex. A {2}, B {2}, C {1} for the indicated read in Fig. 1b-iii), where adjacent identical contig hits are collapsed, and the numbers of collapsed hits are retained. The contig hits are filtered to remove any contigs less than the minimum contig size (*z*, default 1000), and any subsumed contigs.

From this list of contig hits, we can infer oriented pairings between the draft contigs. As well as the obvious adjacent pairings (Ex. A -> B, B -> C in Fig. 1b-iii), we also add pairs based on transitive pairings in the run of contig hits (Ex. A -> C). All transitive pairings are added for lists of contig hits up to *f* (default 10). To avoid the computational overhead becoming too large with many contig hits, when there are more than *f* contigs in a list, transitive pairings are only generated over weakly supported contigs, which are defined as contigs with only a single minimizer hit from the long read (Ex. “C” in the indicated long read in Fig. 1b-iii).

Each of these inferred contig pairs are also oriented relative to each other, and the gap size between them estimated. Each join suggested by a long read *lr* is due to minimizer mappings to the contigs in the pair. Therefore, for each of these joins, there are terminal minimizer hits (mA, mB) for each contig (cA, cB) in a pair, where mA is the last minimizer in contig cA that the long read *lr* hits to, and mB is the first minimizer in contig cB that the long read *lr* matches (Additional File 1: Fig. S1). We compare the strands of canonical minimizers mA and mB in the long read *lr* sketch and the contigs (cA, cB) to orient the assembly sequences. If, for example, mA has the same strand in contig cA and long read *lr*, contig cA is assigned the positive orientation, otherwise, it is reverse-complemented. To estimate the gap size, a similar approach to that employed in ntJoin[25] is used, where the distance between the minimizers mA and mB on the long read *lr* is determined and then corrected for the distance of the minimizers to the contig ends (Additional File 1: Fig. S1).

Finally, after the contig pairs are fully tallied using all of the minimizer sketches from the long reads, a scaffold graph is created, where the nodes are oriented contigs, and directed edges are created between the tallied contig pairs (Fig. 1b-iv). The edge properties are the number of long reads that support the contig pair, and the median estimated gap size. Each contig node is represented in its forward and reverse orientations in the graph, so each tallied pair will be represented twice (ex. A+ → B+, B-→ A- for Fig. 1b-iv).

The scaffold graph is then input to abyss-scaffold[30], a scaffolding layout tool from our ABySS[31] suite of tools, which manipulates and traverses the graph to generate the final scaffold sequences (Fig. 1b-v).

### Test Runs

To test the correction and scaffolding utilities of LongStitch on human data, we obtained ONT long-read data and Illumina HiSeq short-read data for three human individuals[32, 33]: NA12878, NA19240, and NA24385 (Additional File 1: Table S1). We assembled the short-read data using ABySS[31], and the long-read data with Shasta[9] to generate six baseline assemblies to improve using LongStitch. The Shasta assemblies were polished using the corresponding long reads with Racon (v1.4.13) [34]. Assembly tool versions and statistics for the baseline assemblies are summarized in Additional File 1: Tables S2-S3.

We improved each of the baseline assemblies using LongStitch (v1.0.0), optimizing the *k* and *w* parameters for ntLink (v1.0.0), with all other parameters kept at the default values (Tigmint v1.2.3, ARCS/ARKS v1.2.2). Each of the baseline assemblies was also scaffolded with LRScaf (v1.10.0), using parameters -mioll 400 -i0.15 -mxel 500 -mxohl 500 -micl 1000[15]. The long reads were mapped to the draft contigs using minimap2[16] prior to LRScaf, as this is a required pre-processing step for the tool.

All assemblies were analyzed using QUAST v5.0.2 (--fast --scaffold-gap-max-size 100000 --large)[35], and the human reference genome GRCh38. To assess the contiguity of the assemblies, we used both the NG50 and NGA50 lengths. While the NG50 statistic describes that at least half of the genome is in pieces at least the NG50 length, the NGA50 metric is similar, but uses alignment blocks instead of sequence lengths for the calculation. Jupiter plots were also generated to visualize the consistency of each assembly with the human reference genome (ng=75, minBundleSize=50000) [36]. All benchmarking tests were run on a DELL server with 128 Intel(R) Xeon(R) CPU E7–8867 v3, 2.50GHz with 2.6 TB RAM.

## Results and Discussion

To demonstrate the performance of LongStitch in improving upon draft assemblies using long reads, we ran the default steps of the correction and scaffolding pipeline (up to the ntLink stage) on six different human assemblies. We also ran the current state-of-the-art long-read scaffolder, LRScaf[15], on the same data (Fig. 2, Additional File 1: Tables S4-S5).

**Fig. 2.**
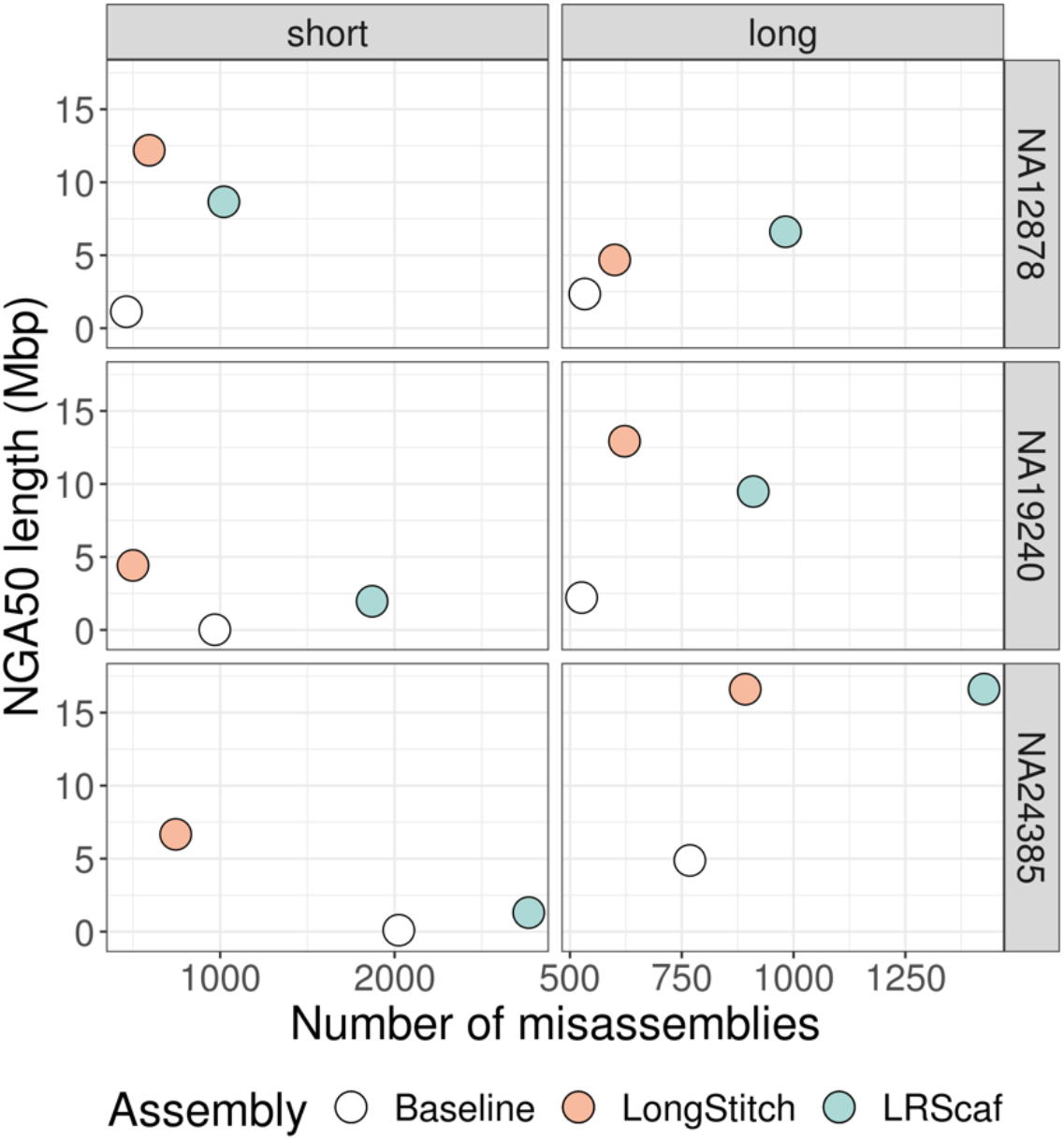
Contiguity and correctness of assemblies improved with LongStitch compared to LRScaf. For three human individuals, NA12878, NA19240 and NA24385, short-read ABySS assemblies and long-read Shasta assemblies (white) were improved using long-read data from the respective individual. The resulting assemblies from LongStitch (orange) and LRScaf (blue) were assessed using QUAST for their contiguity and correctness. For the correction and scaffolding runs shown, the default steps of LongStitch were run (up to the ntLink stage). Ideal assemblies are located in the top-left corner.

For each short-read ABySS assembly, LongStitch significantly improved the contiguity of the baseline assemblies, from a 10.8-fold increase in NGA50 length (1.1 Mbp to 12.2 Mbp) for NA12878 up to a very substantial 304.6-fold increase (14.5 kb to 4.4 Mbp) in NGA50 length for the NA19240 individual. Despite the baseline short-read assembly for the NA19240 individual having the lowest contiguity of the short-read assemblies (NGA50 length = 14.5kb), the LongStitch pipeline was still able to leverage the long-read information to increase the contiguity of this very fragmented assembly to the megabase scale (NGA50 length = 4.4Mbp). This demonstrates a very promising option for a hybrid assembly approach, as scaffolding a fragmented short-read assembly with long reads can generate a very contiguous assembly with an extremely high base accuracy. LongStitch also generates assemblies that are more contiguous than those produced by the LRScaf scaffolder, with final NGA50 lengths up to 5.1-fold higher (1.4-fold, 2.3-fold and 5.1-fold for NA12878, NA19240 and NA24385, respectively). Furthermore, the LongStitch assemblies are more correct, as assessed using QUAST, with LRScaf generating 72.2%, 274.2% and 271.7% more misassemblies than LongStitch for NA12878, NA19240 and NA24385, respectively. In fact, for two of the individuals (NA19240 and NA24385) the assemblies produced by LongStitch have fewer (469 and 1,279, respectively) misassemblies than the baseline, demonstrating the effectiveness of the Tigmint-long misassembly correction step.

As well as showing great potential in a hybrid assembly use-case, the LongStitch pipeline can also utilize long reads to improve Shasta assemblies of the same data. LongStitch improves upon the baseline long-read assembly NGA50 lengths 2.0, 5.8 and 3.4-fold for the NA12878, NA19240 and NA24385 individuals, respectively. This demonstrates that there is still underutilized long-range information in the long-read sequencing data that can be leveraged to improve the assemblies, even after the initial *de novo* Shasta assembly. Compared to LRScaf, LongStitch achieves higher or equivalent NGA50 lengths for NA19240 and NA24385 (1.4-fold higher and equivalent NGA50 lengths, respectively), but does produce a slightly lower NGA50 length for the NA12878 individual. However, similar to the short-read assembly runs, the LongStitch scaffolds have substantially fewer QUAST misassemblies compared to the LRScaf scaffolds, with LongStitch only increasing the misassemblies 12.6 – 18.3% compared to the long-read baselines, whereas LRScaf increased the misassemblies 73.0 – 85.7%. This shows that the LongStitch pipeline, with the combined correction and scaffolding steps, greatly improves the contiguity of the long-read baseline assemblies, and does so with high accuracy, a characteristic that is extremely important to downstream genomics analyses including gene annotation tasks, for example.

We inspected the genome assembly consistency between the LongStitch assemblies and the human reference genome (GRCh38) visually using Jupiter plots[36]. These circos-based[37] representations show sequence alignments between assemblies and the reference genome as coloured bands, and any large-scale misassemblies are immediately evident as interrupting ribbons. Comparing the Jupiter plots generated using the LongStitch and LRScaf assemblies for each of the runs, there are fewer interrupting ribbons in the LongStitch plots in each case, indicating that LongStitch produces fewer large-scale misassemblies (Additional File 1: Fig. S2). This correctness is particularly evident in the runs improving the NA19240 and NA24385 short-read ABySS draft assemblies, where the Tigmint-long correction step was very important to both breaking misassemblies and enabling correct scaffolding with ntLink.

Comparing the benchmarking performances of the tools, LongStitch, despite being a multi-tool pipeline (two tools by default), runs faster than LRScaf for five of the six test runs, with all LongStitch tests running between 3.0 – 4.5 hours (Fig. 3). While LRScaf was 40 minutes faster for the NA12878 short-read assembly test, its runtimes varied quite widely from 2.7h to 44.6h across all runs. Therefore, whereas LRScaf can have unpredictably longer runtimes, the consistent runtimes of LongStitch for a given genome size and read coverage, independent of assembly contiguity, will be very useful when applying the tool to new and larger genomes. While LongStitch did use less memory than LRScaf in five of six tests runs, all runs for both tools used 17.1-23.5 GB of RAM.

**Fig. 3.**
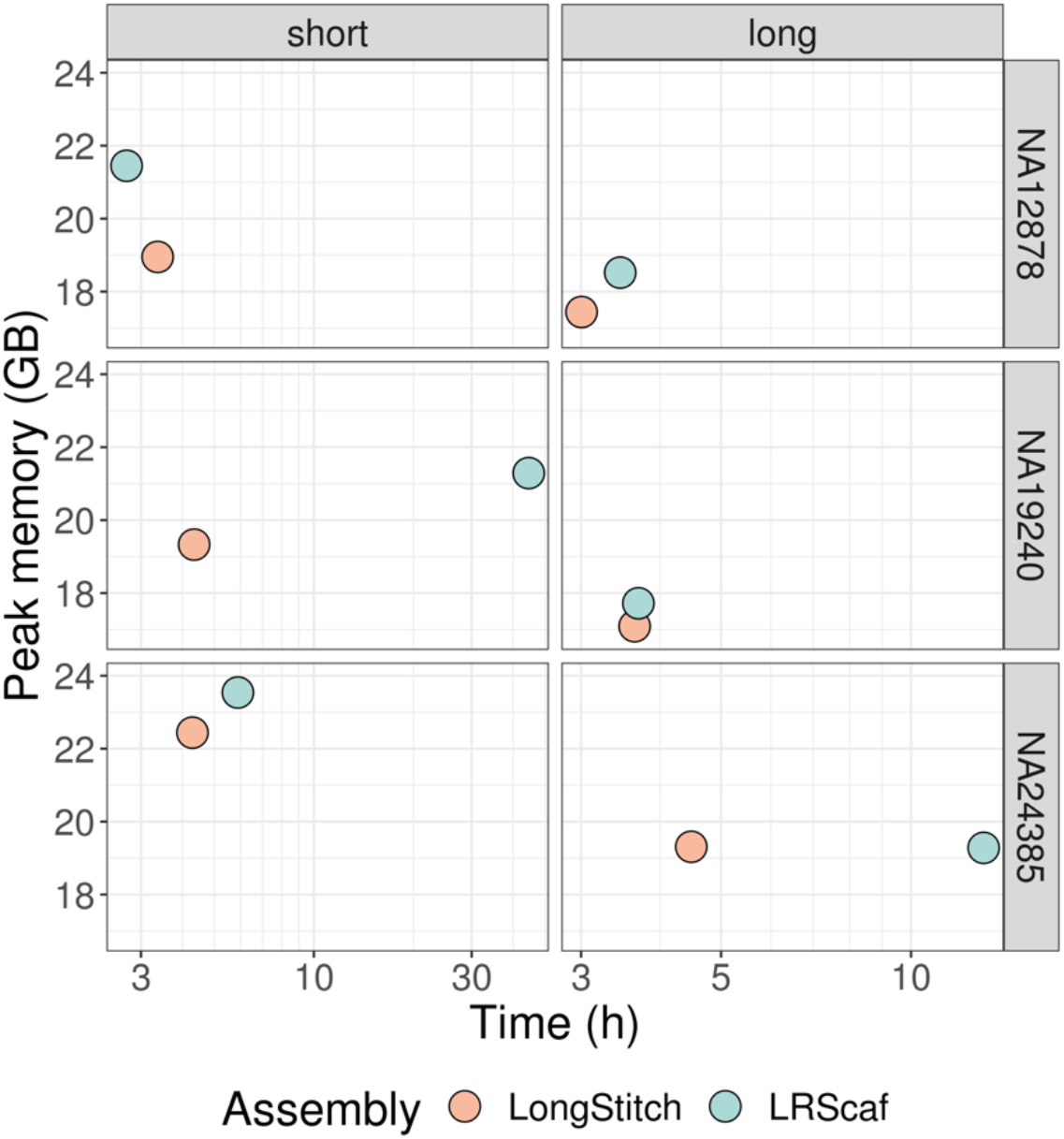
Benchmarking results of improving assemblies with LongStitch or LRScaf. For three human individuals, NA12878, NA19240 and NA24385, the wall-clock time and peak memory is shown for improving short-read ABySS assemblies and long-read Shasta assemblies using the default steps of LongStitch (up to ntLink) (orange) or LRScaf (blue). The peak memory (in gigabytes) is shown on a linear scale, and the wall-clock time (in hours) is shown on a log scale.

While the default mode of the LongStitch pipeline runs two steps: Tigmint-long for misassembly correction, then ntLink for assembly scaffolding with long reads, users can optionally run an extra scaffolding step with ARKS-long to maximize the contiguity improvements. In all cases, this extra step of scaffolding improves the NGA50 metric, showing that this step makes additional correct joins. The improvements in contiguity with ARKS-long are most substantial for the short-read assembly runs (Additional File 1: Fig. S3, Table S6). However, running the extra scaffolding step of course increases the runtime of the whole pipeline and also introduces additional misassemblies (Additional File 1: Fig. S3-S5). Therefore, running the default two-step LongStitch pipeline or additionally running the ARKS-long step is a decision open to the user, and will likely depend on the user’s particular use case and application.

We previously demonstrated the effectiveness of using minimizers for genome scaffolding with our reference-guided scaffolder, ntJoin[25], and we find that minimizers also exhibit great utility in fast and accurate mapping of long reads to assemblies using ntLink. The *k* and *w* parameters of ntLink do impact the resulting scaffolding, but we find that ntLink works well over a range of these values (Additional File 1: Fig. S6, S7). Furthermore, the NG50 metrics of the resulting assemblies, which are calculated without using a reference, show a similar pattern to the NGA50 reference-based contiguity metric, demonstrating that a user can optimize these *k* and *w* parameters without the requirement of a reference. As well as ordering and orienting contigs, ntLink uses the minimizer mappings of the long reads to the draft assembly to estimate the gap sizes in the output scaffolds. In the current implementation, the ntLink code creates the scaffold graph, which is traversed by abyss-scaffold[30], a scaffold layout algorithm from our ABySS suite of tools, but the tool is flexible to using other scaffold layout algorithms if desired, similar to our ARKS scaffolding tool.

## Conclusions

We have demonstrated the use of our scalable long-read correction and scaffolding pipeline, LongStitch, on a variety of human datasets and assemblies, and show that it runs efficiently and generates high-quality final assemblies. When embarking on *de novo* assembly projects, different groups will have varying combinations of sequencing data, and it is important to have tools available that are useful for a range of use cases. With the LongStitch pipeline, long reads are used to improve upon an input draft assembly from any data type. Therefore, if a project solely uses long reads, the LongStitch pipeline is able to further improve upon *de novo* long-read assemblies. However, if the baseline assembly is a short-read assembly, a linked-read assembly or even an assembly incorporating multiple data types, LongStitch is also valuable for facilitating additional improvements. Due to its efficiency and flexibility to many different *de novo* genome assembly projects, we expect LongStitch to be widely beneficial to the research community.

## Availability and Requirements

**Project name:** LongStitch

**Project home page:** https://github.com/bcgsc/longstitch

**Operating system(s):** Platform independent

**Programming language:** Python, C++, GNU Make

**Other requirements:** Tigmint, ntLink, ARCS/ARKS, ABySS

**License:** GPL v3

**Any restrictions to use by non-academics:** No

## Declarations

### Availability of data and materials

The sources of all data used and/or analyzed in this study are listed in Additional File 1: Tables S1 and S3.

### Authors’ contributions

LC, RLW and IB were major contributors to the ideas for the LongStitch pipeline and its steps. LC designed and coded ntLink, ran the benchmarking tests, analyzed the results and wrote the manuscript. JXL designed and implemented Tigmint-long, and TL designed and implemented ARKS-long. JW contributed and optimized the code for pseudo-linked read generation for Tigmint-long and ARKS-long. VN contributed and optimized the code for minimizer generation. All authors contributed to editing the manuscript, and all authors read and approved the final manuscript.

## Acknowledgements

We thank Justin Chu for his valuable contributions to brainstorming discussions early in the project, which helped drive the development of the algorithms.

