## Additional File 1 for "LongStitch: High-quality genome assembly correction and scaffolding using long reads"

### Supplementary Tables

**Table S1. Sequencing data used for the correction and scaffolding runs.**

| Individual | Reads type | Fold coverage | Platform | Accession(s)/Source |
|---|---|---|---|---|
| NA12878 | Short | 54 | Illumina HiSeq | Illumina basespace – run title: "HiSeq 2500: TruSeq PCR-Free DNA 2x251 (NA12878)" |
| NA12878 | MPET | N/A | Illumina HiSeq | ERR262997 |
| NA12878 | Long | 39 | ONT | SRR10965087 |
| NA19240 | Short | 62 | Illumina HiSeq | SRR3189758, SRR3189759 |
| NA19240 | Long | 49 | ONT | ERR3219854, ERR3219857 |
| NA24385 | Short | 58 | Illumina HiSeq | SRR11321732 |
| NA24385 | Long | 51 | ONT | s3://ont-open-data/gm24385_2020.11/analysis/r9.4.1/20201026_1644_2-E5-H5_PAG07162_d7f262d5/guppy_v4.0.11_r9.4.1_hac_prom/basecalls.fastq.gz |

ONT: Oxford Nanopore Technologies

**Table S2. Baseline assemblies used for correction and scaffolding runs.** 'n' is the number of contigs over 3kb. Default parameters were used for the assemblers, with the exception of the parameters listed.

| Individual | Assembly reads | Assembler | Parameters | n | NG50 (bp) | NGA50 (bp) | Largest contig (bp) | Number of misassemblies |
|---|---|---|---|---|---|---|---|---|
| NA12878 | Short + MPET | ABySS 2.1.5 | k=128, kc=3, B=115, j=24 | 6,656 | 1,175,297 | 1,131,267 | 7,807,150 | 462 |
| NA12878 | Long | Shasta 0.5.1 | --threads 48 | 3,596 | 2,617,486 | 2,346,570 | 17,057,497 | 533 |
| NA19240 | Short | ABySS 2.2.3 | k=112, kc=3, B=150, j=48 | 190,813 | 14,618 | 14,512 | 202,526 | 969 |
| NA19240 | Long | Shasta 0.5.1 | --threads 48 | 4,316 | 2,389,148 | 2,213,238 | 13,988,698 | 526 |
| NA24385 | Short | ABySS 2.2.3 | k=112, kc=3, B=150, j=48 | 44,555 | 102,102 | 97,340 | 1,078,380 | 2,024 |
| NA24385 | Long | Shasta 0.5.1 | --threads 48 | 4,268 | 5,734,421 | 4,876,524 | 30,135,269 | 768 |

**Table S3. Availability of baseline assemblies used for correction and scaffolding runs.**

| Individual | Assembly reads | Assembler | URL |
|---|---|---|---|
| NA12878 | Short + MPET | ABySS 2.1.5 | https://www.bcgsc.ca/downloads/btl/LongStitch/baseline_assemblies/NA12878/abyss/NA12878.abyss-mpet.fa.gz |
| NA12878 | Long | Shasta 0.5.1 | https://www.bcgsc.ca/downloads/btl/LongStitch/baseline_assemblies/NA12878/shasta/NA12878.shasta-polished.fa.gz |
| NA19240 | Short | ABySS 2.2.3 | https://www.bcgsc.ca/downloads/btl/LongStitch/baseline_assemblies/NA19240/abyss/NA19240.abyss.fa.gz |
| NA19240 | Long | Shasta 0.5.1 | https://www.bcgsc.ca/downloads/btl/LongStitch/baseline_assemblies/NA19240/shasta/NA19240.shasta-polished.fa.gz |
| NA24385 | Short | ABySS 2.2.3 | https://www.bcgsc.ca/downloads/btl/LongStitch/baseline_assemblies/NA24385/abyss/NA24385.abyss.fa.gz |
| NA24385 | Long | Shasta 0.5.1 | https://www.bcgsc.ca/downloads/btl/LongStitch/baseline_assemblies/NA24385/shasta/NA24385.shasta-polished.fa.gz |

**Table S4. Contiguity, correctness and benchmarking statistics for running the default steps of LongStitch (up to ntLink).** Default parameters were used, with the exception of the ntLink parameters listed. Misassemblies were assessed using QUAST. 'n' is the number of contigs over 3kb.

| Individual | Assembler | ntLink Parameters | n | NG50 (Mbp) | NGA50 (Mbp) | Number of misassemblies | Time (h) | Peak memory (GB) |
|---|---|---|---|---|---|---|---|---|
| NA12878 | ABySS | k=32, w=100 | 2,521 | 15.32 | 12.19 | 593 | 3.37 | 18.95 |
| NA12878 | Shasta | k=24, w=250 | 3,126 | 5.70 | 4.68 | 600 | 3.00 | 17.44 |
| NA19240 | ABySS | k=40, w=100 | 5,437 | 5.11 | 4.42 | 500 | 4.34 | 19.33 |
| NA19240 | Shasta | k=24, w=250 | 2,571 | 18.66 | 12.93 | 622 | 3.65 | 17.09 |
| NA24385 | ABySS | k=40, w=500 | 5,375 | 8.12 | 6.66 | 745 | 4.29 | 22.44 |
| NA24385 | Shasta | k=40, w=500 | 3,440 | 26.97 | 16.61 | 892 | 4.48 | 19.31 |

**Table S5. Contiguity, correctness and benchmarking statistics for LRScaf.** Misassemblies were assessed using QUAST. 'n' is the number of contigs over 3kb.

| Individual | Assembler | n | NG50 (Mbp) | NGA50 (Mbp) | Number of misassemblies | Time (h) | Peak memory (GB) |
|---|---|---|---|---|---|---|---|
| NA12878 | ABySS | 2,525 | 11.49 | 8.66 | 1,021 | 2.71 | 21.45 |
| NA12878 | Shasta | 1,711 | 8.51 | 6.61 | 982 | 3.46 | 18.52 |
| NA19240 | ABySS | 3,079 | 3.10 | 1.96 | 1,871 | 44.61 | 21.29 |
| NA19240 | Shasta | 2,068 | 12.40 | 9.48 | 910 | 3.70 | 17.72 |
| NA24385 | ABySS | 4,015 | 2.23 | 1.30 | 2,769 | 5.89 | 23.54 |
| NA24385 | Shasta | 3,030 | 24.28 | 16.61 | 1,426 | 13.03 | 19.28 |

**Table S6. Contiguity, correctness and benchmarking statistics for LongStitch including the optional ARKS-long step.** Default parameters were used, with the exception of the ntLink parameters listed. Misassemblies were assessed using QUAST. 'n' is the number of contigs over 3kb.

| Individual | Assembly | ntLink Parameters | n | NG50 (Mbp) | NGA50 (Mbp) | Number of misassemblies | Time (h) | Peak memory (GB) |
|---|---|---|---|---|---|---|---|---|
| NA12878 | ABySS | k=32, w=100 | 2,135 | 20.89 | 14.12 | 746 | 6.76 | 18.95 |
| NA12878 | Shasta | k=24, w=250 | 2,702 | 6.27 | 5.08 | 752 | 6.46 | 17.44 |
| NA19240 | ABySS | k=40, w=100 | 4,359 | 10.94 | 6.94 | 626 | 9.07 | 19.33 |
| NA19240 | Shasta | k=24, w=250 | 2,120 | 23.38 | 13.71 | 773 | 8.13 | 17.09 |
| NA24385 | ABySS | k=40, w=500 | 4,453 | 13.88 | 9.15 | 1,003 | 9.59 | 22.44 |
| NA24385 | Shasta | k=40, w=500 | 3,049 | 32.20 | 16.66 | 969 | 9.70 | 19.31 |

#### Supplementary Figures

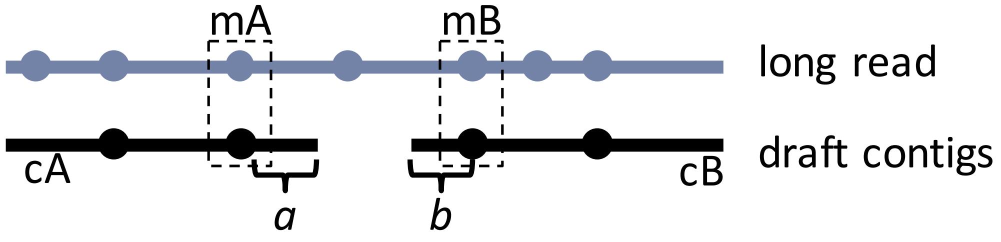

$$gap\ size = |pos(mA, long\_read) - pos(mB, long\_read)| - a - b$$
where $a = distance\ from\ pos(mA, draft)\ to\ end\ of\ cA$
and $b = distance\ from\ pos(mB, draft)\ to\ end\ of\ cB$

**Figure S1. Schematic describing the gap size estimation algorithm in ntLink.**

**Figure S2. Jupiter consistency plots showing the contiguity and correctness of LongStitch and LRScaf assemblies.** Each (A) short-read ABySS assembly and (B) long-read Shasta assembly was improved with LongStitch (default steps) or LRScaf, and aligned to the human reference genome (GRCh38). The alignments are plotted as coloured ribbons in each Jupiter plot, with large-scale misassemblies evident as interrupting ribbons.

| **(A)** | **short** | |
|---|---|---|
| **Individual** | **LongStitch** | **LRScaf** |
| **NA12878** | 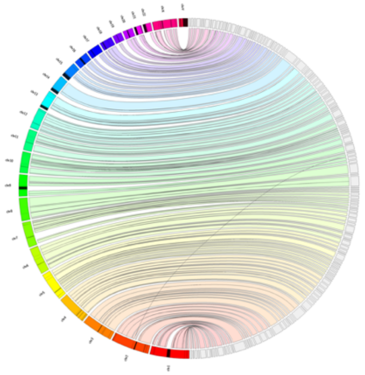 | 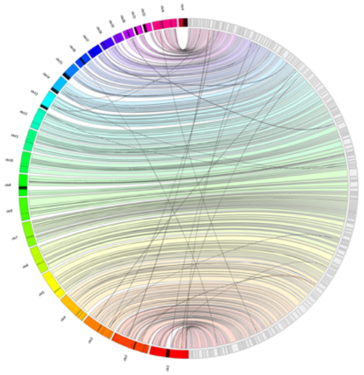 |
| **NA19240** | 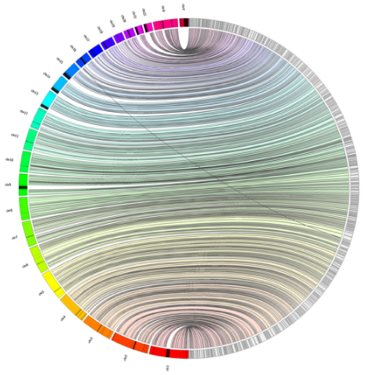 | 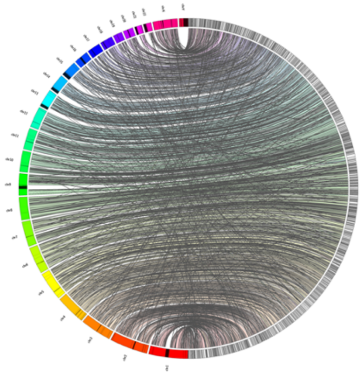 |
| **NA24385** | 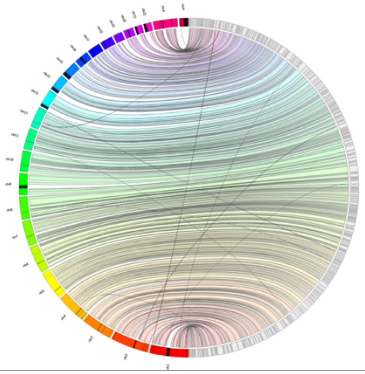 | 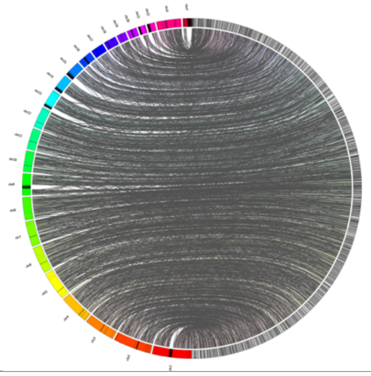 |

| (B) | long | |
|---|---|---|
| Individual | LongStitch | LRScaf |
| NA12878 | 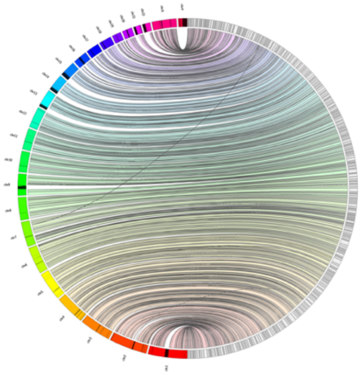 | 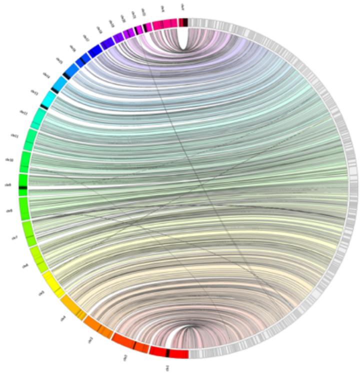 |
| NA19240 | 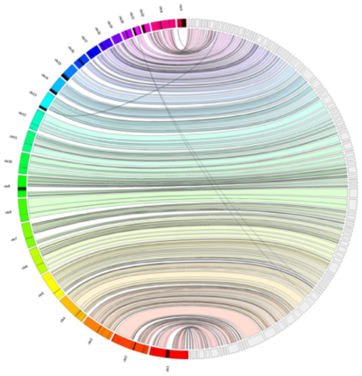 | 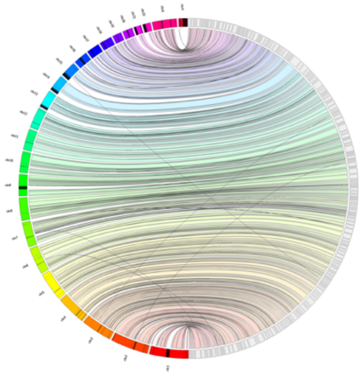 |
| NA24385 | 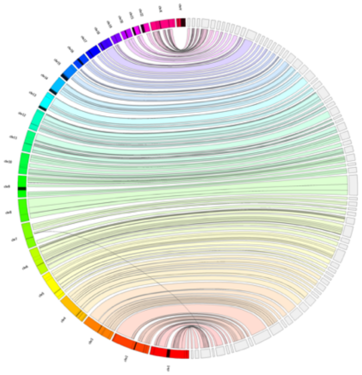 | 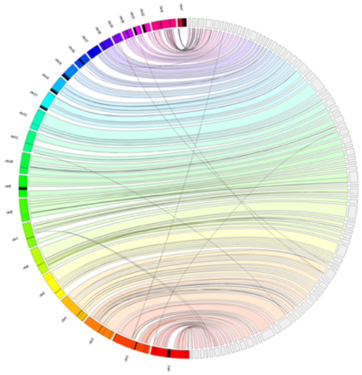 |

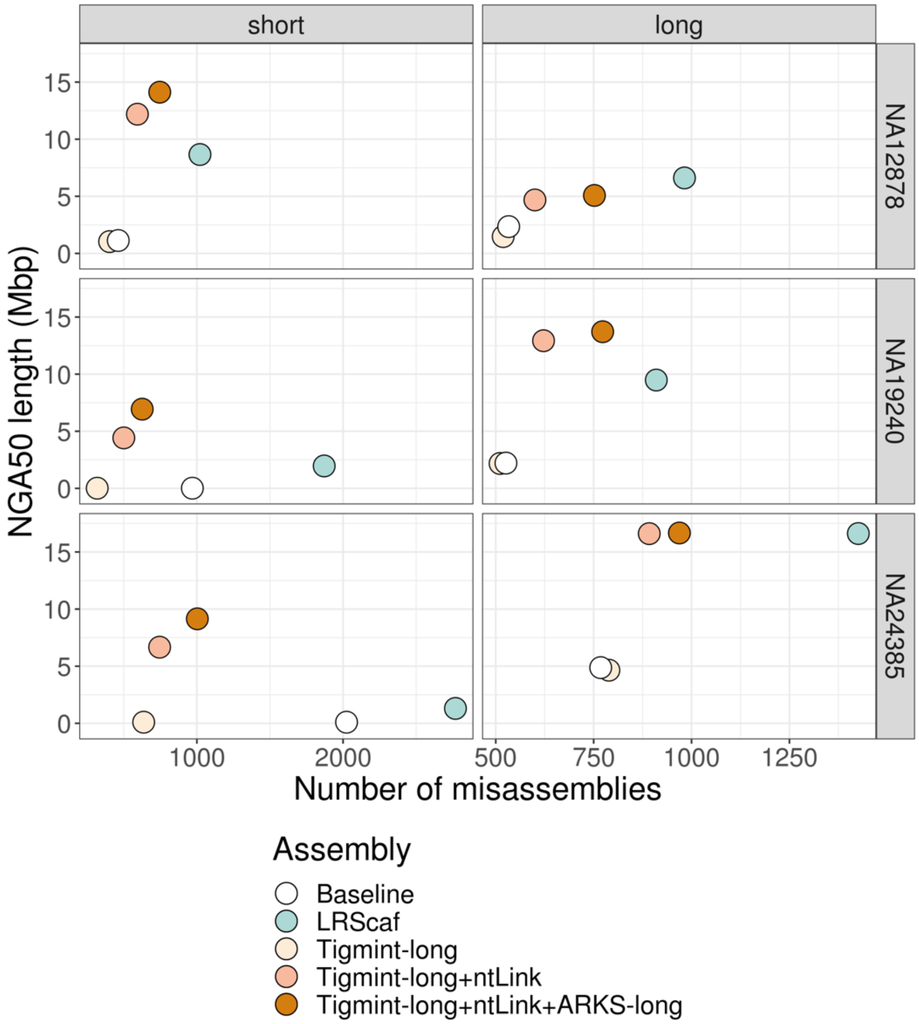

**Figure S3. Contiguity and correctness results for each step the LongStitch pipeline, including the optional ARKS-long step, and LRScaf.** Each baseline assembly (white) was improved using LongStitch (orange) and LRScaf (blue).

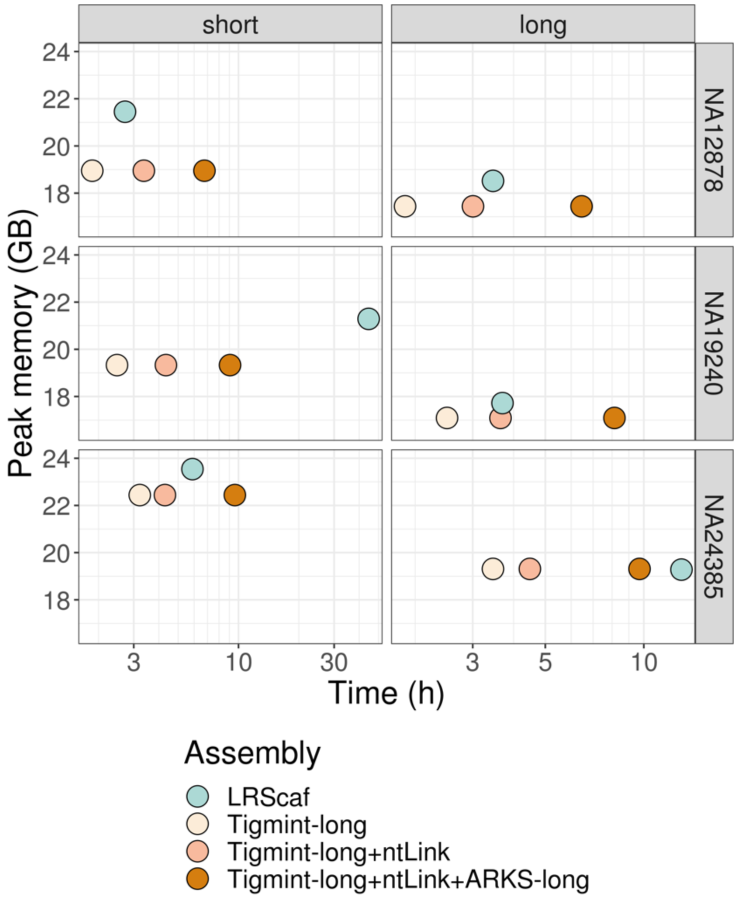

**Figure S4. Benchmarking results for LRScaf and the LongStitch pipeline including the optional ARKS-long step.**

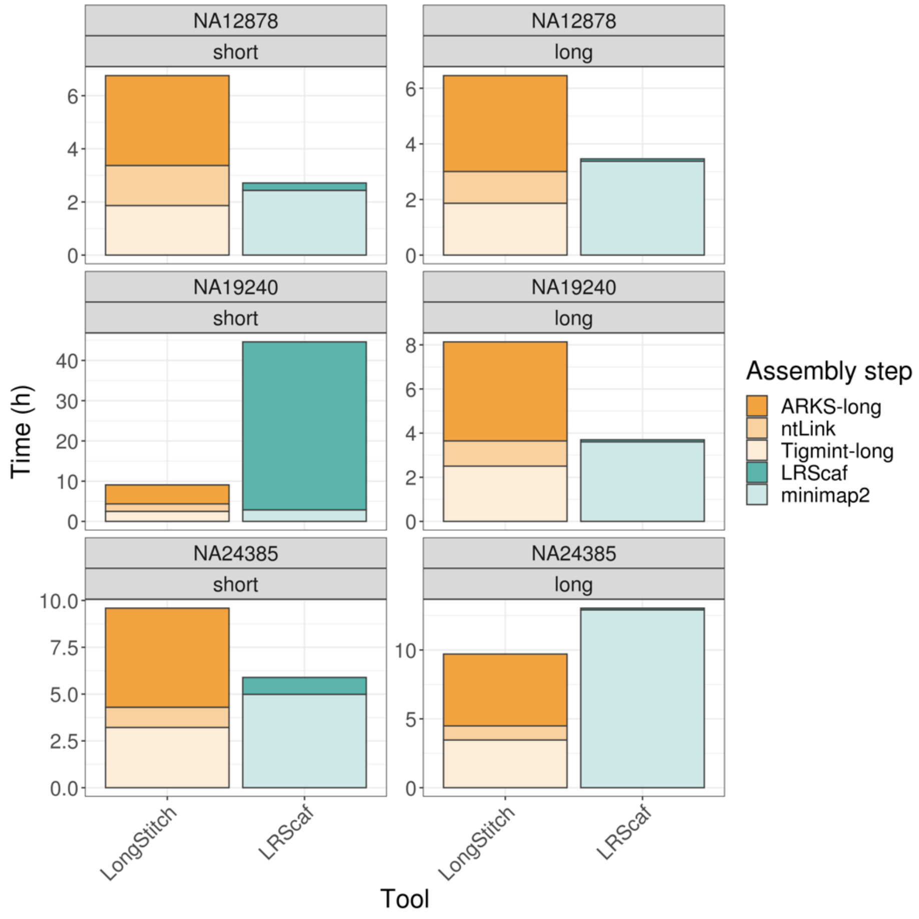

**Figure S5. Time breakdown for each of the steps in LongStitch, including the optional ARKS-long step, and LRScaf.**

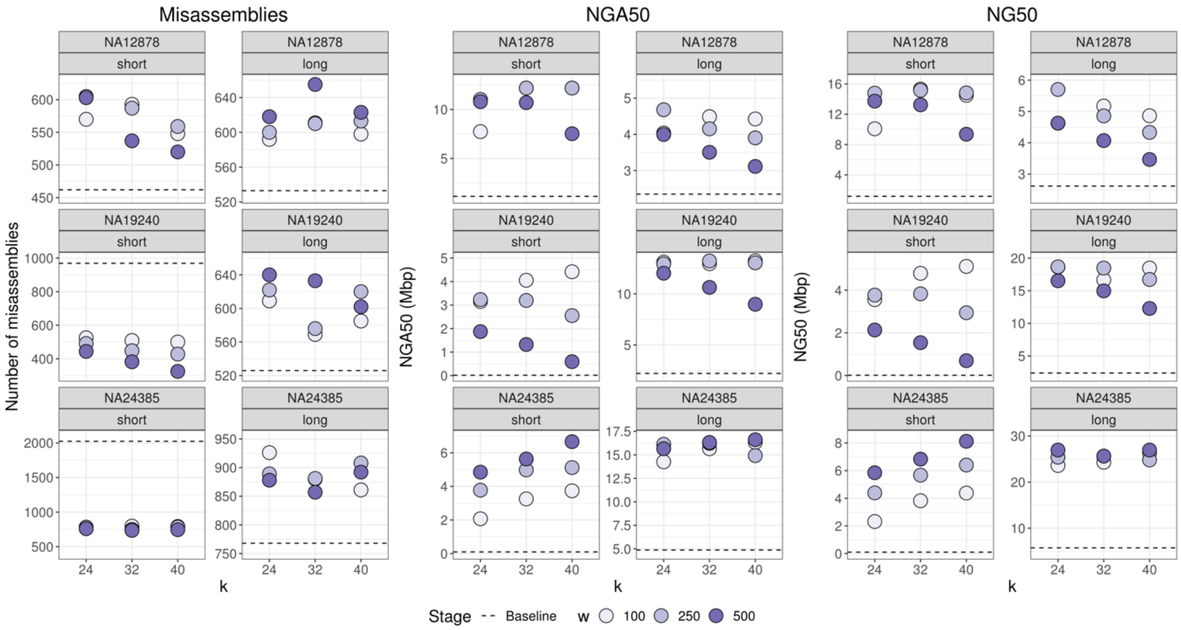

**Figure S6. Contiguity and correctness results from sweeping on the ntLink *k* and *w* parameters for the default steps of LongStitch.** The baseline assembly statistics are indicated with a horizontal dashed line.

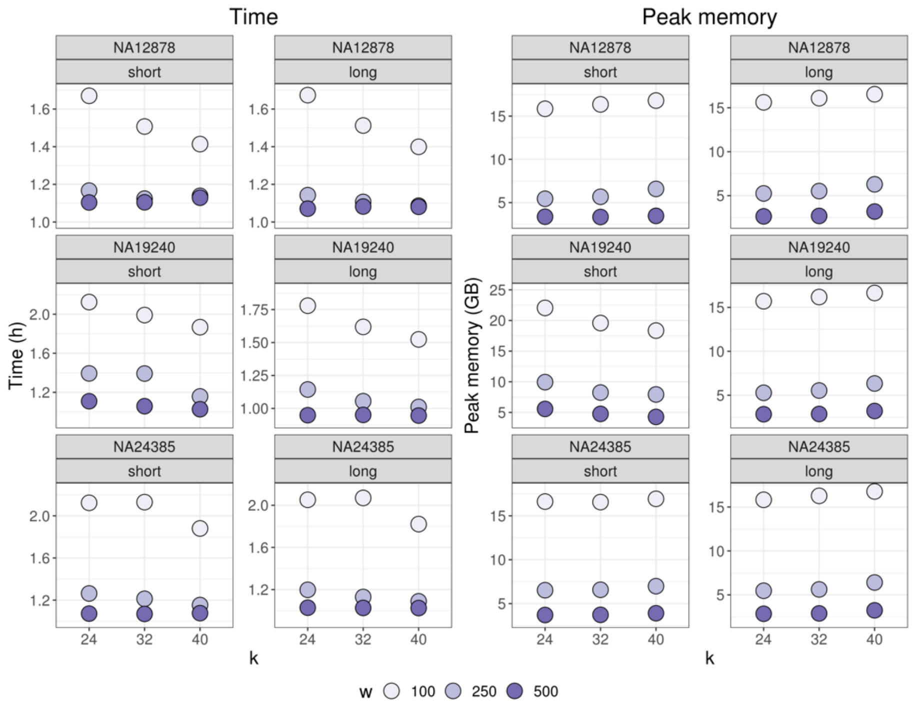

**Figure S7. Benchmarking results from sweeping on the ntLink *k* and *w* parameters for the default steps of LongStitch.**
